# Role of Inflammasome-independent Activation of IL-1β by the *Pseudomonas aeruginosa* Protease LasB

**DOI:** 10.1101/2020.05.18.101303

**Authors:** Josh Sun, Doris L. LaRock, Elaine A. Skowronski, Jacqueline M. Kimmey, Joshua Olson, Zhenze Jiang, Anthony J. O’Donoghue, Victor Nizet, Christopher N. LaRock

## Abstract

Pulmonary damage by *Pseudomonas aeruginosa* during cystic fibrosis lung infection and ventilator-associated pneumonia is mediated both by pathogen virulence factors and host inflammation. Impaired immune function due to tissue damage and inflammation, coupled with pathogen multidrug resistance, complicates management of these deep-seated infections. Therefore, preservation of lung function and effective immune clearance may be enhanced by selectively controlling inflammation. Pathological inflammation during *P. aeruginosa* pneumonia is driven by interleukin-1β (IL-1β). This proinflammatory cytokine is canonically regulated by caspase-family inflammasome proteases, but we report that plasticity in IL-1β proteolytic activation allows for its direct maturation by the pseudomonal protease LasB. LasB promotes IL-1β activation, neutrophilic inflammation, and destruction of lung architecture characteristic of severe *P. aeruginosa* pulmonary infection. Discovery of this IL-1β regulatory mechanism provides a distinct target for anti-inflammatory therapeutics, such that matrix metalloprotease inhibitors blocking LasB limit inflammation and pathology during *P. aeruginosa* pulmonary infections.

**Highlights:**

- IL-1β drives pathology during pulmonary infection by *Pseudomonas aeruginosa*.
- The *Pseudomonas* protease LasB cleaves and activates IL-1β independent of canonical and noncanonical inflammasomes
- Metalloprotease inhibitors active against LasB limit inflammation and bacterial growth

**Research in Context:** Inflammation is highly damaging during lung infections by the opportunistic pathogen *Pseudomonas aeruginosa*. Sun et al. demonstrate that the *Pseudomonas* LasB protease directly activates IL-1β in an inflammasome-independent manner. Inhibition of IL-1β conversion by LasB protects against neutrophilic inflammation and destruction of the lung. Adjunctive therapeutics that limit pathological inflammation induced by infection would be beneficial for the treatment of pulmonary infections when used with conventional antibiotics.

## Introduction

*Pseudomonas aeruginosa* is a prominent cause of severe opportunistic pulmonary infections associated with mechanical ventilation and the genetic disease cystic fibrosis (CF). *P. aeruginosa* infection is often refractory to antibiotic therapy due to multidrug resistance, making it a World Health Organization and U.S. Centers for Disease Control priority pathogen for therapeutic development. *P. aeruginosa* infection destroys lung architecture and function due to inflammatory- and neutrophil-mediated degradation of mucin layers and structural proteins of the pulmonary connective tissue ^1,2^. Neutrophil cytokines such as IL-1β ^3,4^ and IL-8 ^5^, the latter itself regulated by IL-1β ^6^, initiate and maintain this inflammatory cycle. Anti-inflammatory agents can mitigate tissue destruction to preserve pulmonary function during *P. aeruginosa* pneumonia ^7^ and CF ^8,9^.

Newly synthesized IL-1β (pro-IL-1β) is inactive and requires proteolytic processing into a mature active form. Canonically, this is carried out by the inflammasome, a macromolecular complex of intracellular pattern recognition receptors and the proteases caspase-1 or caspase-11 ^10^. During infection, inflammasomes are formed upon detection of pathogen-associated molecular patterns (PAMPs), including many present in *P. aeruginosa* such as flagellin (FliC), the type III secretion basal body rod (PscI), the type IV pilin (PilA), RhsT, exolysin (ExlA), exotoxin A (ExoA), cyclic 3′-5′ diguanylate (c-di-GMP), and lipopolysaccharide (LPS), which are varyingly detected by NLRC4, NLRP3, or caspase-11 ^11-19^. Some pathogens limit inflammation by targeting the inflammasome ^20^, and *P. aeruginosa* dampens inflammasome activation via the effector ExoU ^18^. Despite the multitude of inflammasome-activating signals that *P. aeruginosa* express, caspases, NLRP3, and NLRC4 are not essential for pro-IL-1β maturation in macrophages, epithelial cells, or neutrophils infected with *P. aeruginosa* ^21,22^. Correspondingly, *P. aeruginosa*-infected caspase-1^-/-^ and caspase-1/11^-/-^ mice succumb to a destructive neutrophilic pulmonary inflammation against which IL-1 receptor (IL-1R1^-/-^) mice are protected ^23^. These observations highlight the contribution of IL-1β to *P. aeruginosa* infection but suggest there are mechanisms for its maturation other than the inflammasome.

The pathological cascade of protease dysregulation and activation seen during severe *P. aeruginosa* lung infections provide a possibility for IL-1β maturation by alternative mechanisms. Caspase-8 ^24-26^, and the neutrophil granular proteases elastase (NE) and proteinase 3 (PR3) ^3,4,27^, cleave pro-IL-1β, but this does not always result in active cytokine ^28^. Bronchial secretions, however, also possess abundant protease activity from microbial sources ^2^. Here we find that IL-1β is not exclusively matured by host proteases, and that *P. aeruginosa* protease LasB also drives this inflammatory pathway. Targeting this bacterial protease may, therefore, provide supportive therapy to limit inflammatory pathology in pulmonary infection.

## Results

### IL-1β drives neutrophilic inflammation during *P. aeruginosa* lung infection

Inflammation drives poor clinical outcomes during *P. aeruginosa* lung infection ^29^. C57Bl/6 mice infected intratracheally with *P. aeruginosa* had markedly disrupted airway architecture within 24 h, concurrent with neutrophil infiltration into the lung tissue and bronchoalveolar lavage fluid (BAL) (Figure 1A). We examined the contribution of pro-inflammatory cytokines to this process using the FDA-approved IL-1 receptor (IL-1R1) antagonist anakinra, which directly inhibits both IL-1β and IL-1α, but not other critical proinflammatory cytokines such as KC/CXCL1, IL-6, or TNFα (Figure 1B). As observed during human infections, *P. aeruginosa* persisted in the BAL (Figure 1C) and lung tissue (Figure 1D) despite significant neutrophil infiltration that was partially IL-1-dependent (Figure 1E).

**Figure 1.**
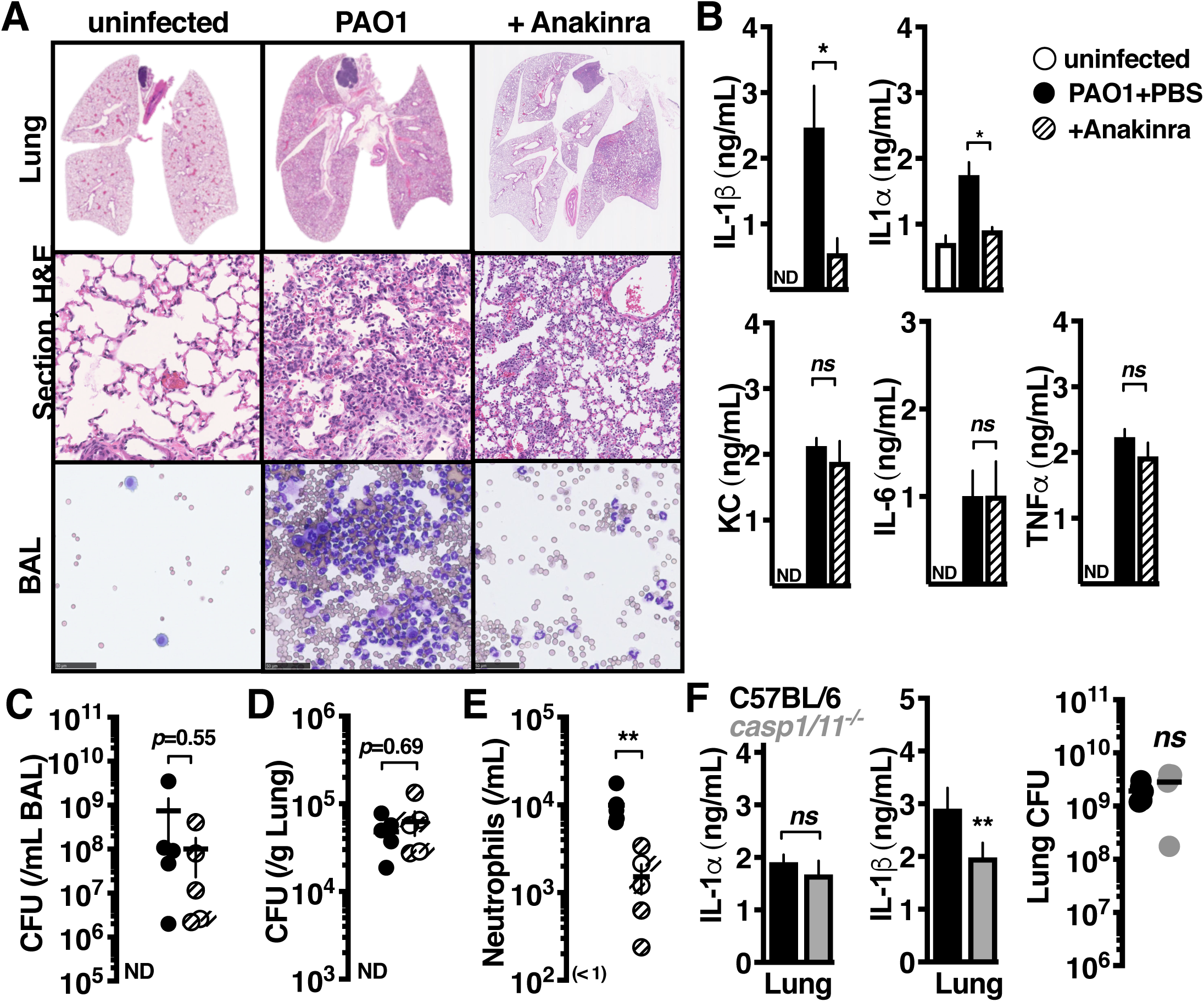
IL-1β drives neutrophilic inflammation during *P. aeruginosa* lung infection. C57BL/6 mice intratracheally infected with 10^7^ colony forming units (CFU) of PAO1 and treated with anakinra (50 µg/kg) or PBS control, compared to uninfected mice. Mice were euthanized after 24 h and (**A**) lung histology sections or cytological smears of bronchoalveolar lavage fluid (BAL) prepared with differential MGG stain, (**B**) BAL cytokines measured by enzyme-linked immunosorbent assay, (**C-D**) bacterial CFU in BAL or lung homogenate, and (**E**) BAL neutrophils enumerated. (**F**) C57BL/6 or isogeneic caspase-1/11^-/-^ mice intratracheally infected with 10^7^ CFU PAO1 24 h, euthanized, and BAL cytokines measured by enzyme-linked immunosorbent assay. Where applicable, data are mean ± SEM and represent at least 3 independent experiments. Significance determined by Mann-Whitney U-test (\**P* < 0.05, \*\**P* < 0.005).

IL-1β is typically released by secretion or cell lysis and requires additional maturation, activities which are all mediated by the inflammasome proteases caspases -1 or -11 ^10^. CFU and release of IL-1α was unaltered in *P. aeruginosa*-infected caspase-1/11^-/-^ C57Bl/6 mice, but surprisingly, IL-1β release was also only modestly attenuated (Figure 1F). This pool of extracellular IL-1β has the potential to mediate proinflammatory signaling as an IL-1R1 agonist when the inhibitory pro-domain has been removed. Neutrophil granular proteases may provide such activation ^3,4,15,21^, however, since neutrophil recruitment is itself IL-1β-dependent (Figure 1A, 1E), and these neutrophils themselves may later inactivate IL-1β ^28^, we reasoned that additional proteases initiate the process.

### *P. aeruginosa* induces IL-1β maturation independent of the inflammasome

To more specifically measure only IL-1β that is active, we made use of transgenic reporter cells expressing luciferase under the control of the IL-1R (Figure 2A) similar to previously ^30^. Consistent with our *in vivo* observations, caspase-1/11^-/-^ bone-marrow-derived macrophages (BMM) still released cytokines that activated IL-1R1 reporter cells upon infection with *P. aeruginosa* PAO1 (Figure 2B). This activity was conserved across numerous *P. aeruginosa* isolates. In contrast, an ionophore that activates the NLRP3 inflammasome, nigericin, was completely dependent on caspases for the activation of IL-1 signaling. Furthermore, *P. aeruginosa* infection of human cell lines relevant to lung infection (macrophages, THP-1; neutrophils, HL60; type II alveolar epithelial cells, A549) still stimulated IL-1 signaling in the presence of the caspase-1/11-specific inhibitor YVAD-cmk (Figure 2C). Monoclonal antibodies specific to IL-1R1 or IL-1β, but not IL-1α, inhibited IL-1 signal from caspase-1/11^-/-^ BMM (Figure 2D). The absolute quantity of each cytokine measured by enzyme-linked immunosorbent assay (pro- and mature-forms) remained unchanged (Figure 2D). Together, these results indicate that *P. aeruginosa* stimulates IL-1 signaling through a pool of extracellular IL-1β that is active and matured independently of caspase-1/11.

**Figure 2.**
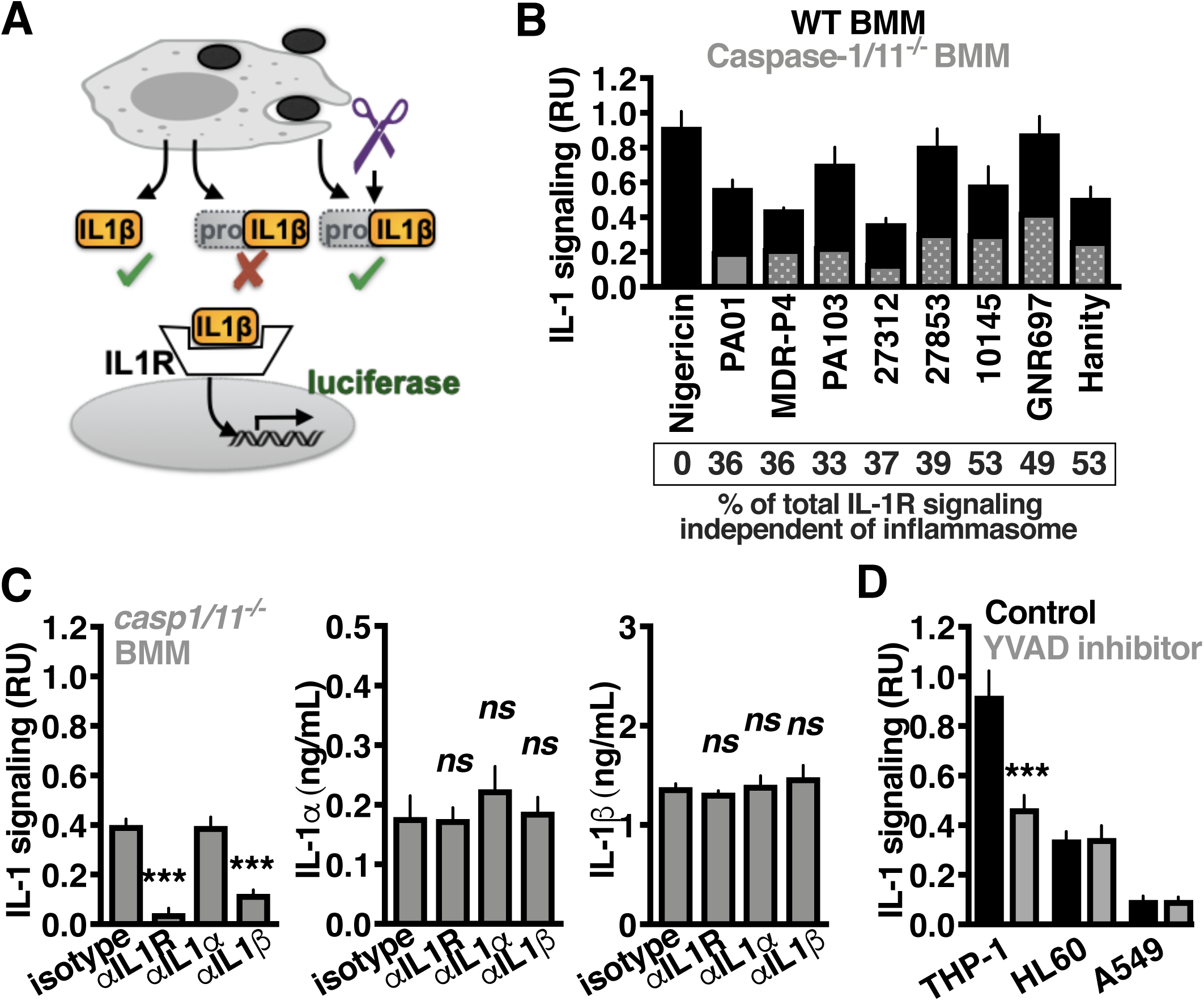
*P. aeruginosa* induces IL-1β maturation independent of the inflammasome. (**A**) Diagram of IL-1 reporter assay. Pro-IL-1β does not induce signaling through the IL-1R. Removal of the pro-domain, intracellularly or extracellularly, by any protease that can do so, results in an active cytokine with proinflammatory activity. (**B**) Relative IL-1 signaling by caspase-1/11^-/-^ (grey) or control C57Bl/6 BMMs (black) after 2 h co-incubation of the indicated *Pseudomonas* strains. Nigericin (5 µM) is included as a positive control for inflammasome-dependent IL-1β maturation. (**C**) Mature IL-1 and enzyme-linked immunosorbent assay measurement of IL-1α and IL-1β present in any form, released from PAO1-infected caspase-1/11^-/-^ BMM with monoclonal antibodies neutralizing IL-1R1, IL-1α, IL-1β, or an isotype control. (**D**) Relative IL-1 signaling by human THP-1 macrophages, HL60 neutrophils, or A549 epithelial cells treated with caspase-1 inhibitor (YVAD) or control (Mock) 1 h prior to infection. Infections were at MOI=10 and after 2 h the supernatant collected and mature IL-1 quantified using IL-1R1 reporter cells. Where applicable, data are mean ± SEM and represent at least 3 independent experiments; significance determined by unpaired two-tailed Student’s T-test, \**P* < 0.05.

### IL-1β is activated by the *P. aeruginosa* LasB protease

Proteases contributing to IL-1β activation were evaluated using small molecule inhibitors specific to each protease class. Inhibition of metalloproteases, and not cysteine proteases (e.g. caspases-1, 11, and 8) or serine proteases (e.g. NE and PR3), abrogated IL-1β signaling in *P. aeruginosa*-infected caspase-1/11^-/-^ BMM (Figure 3A). *P. aeruginosa* encodes several secreted metalloproteases, and by examining mutants of each (Δ*lasA*, Δ*lasB*, Δ*aprA*), we found LasB to be the most active protease overall as measured by hydrolysis of casein during agar plate growth (Figure 3B), and was the major contributor to caspase-1/11-independent IL-1β signaling (Figure 3C). Complementation with the LasB coding sequence under its native promoter restored the ability of Δ*lasB P. aeruginosa* to induce IL-1β signaling in infected caspase-1/11^-/-^ BMM (Figure 3D). Furthermore, activation was independent of *il1b* expression (Figure 2E). These data show that LasB induces IL-1 signaling independently of caspase-1/11.

**Figure 3.**
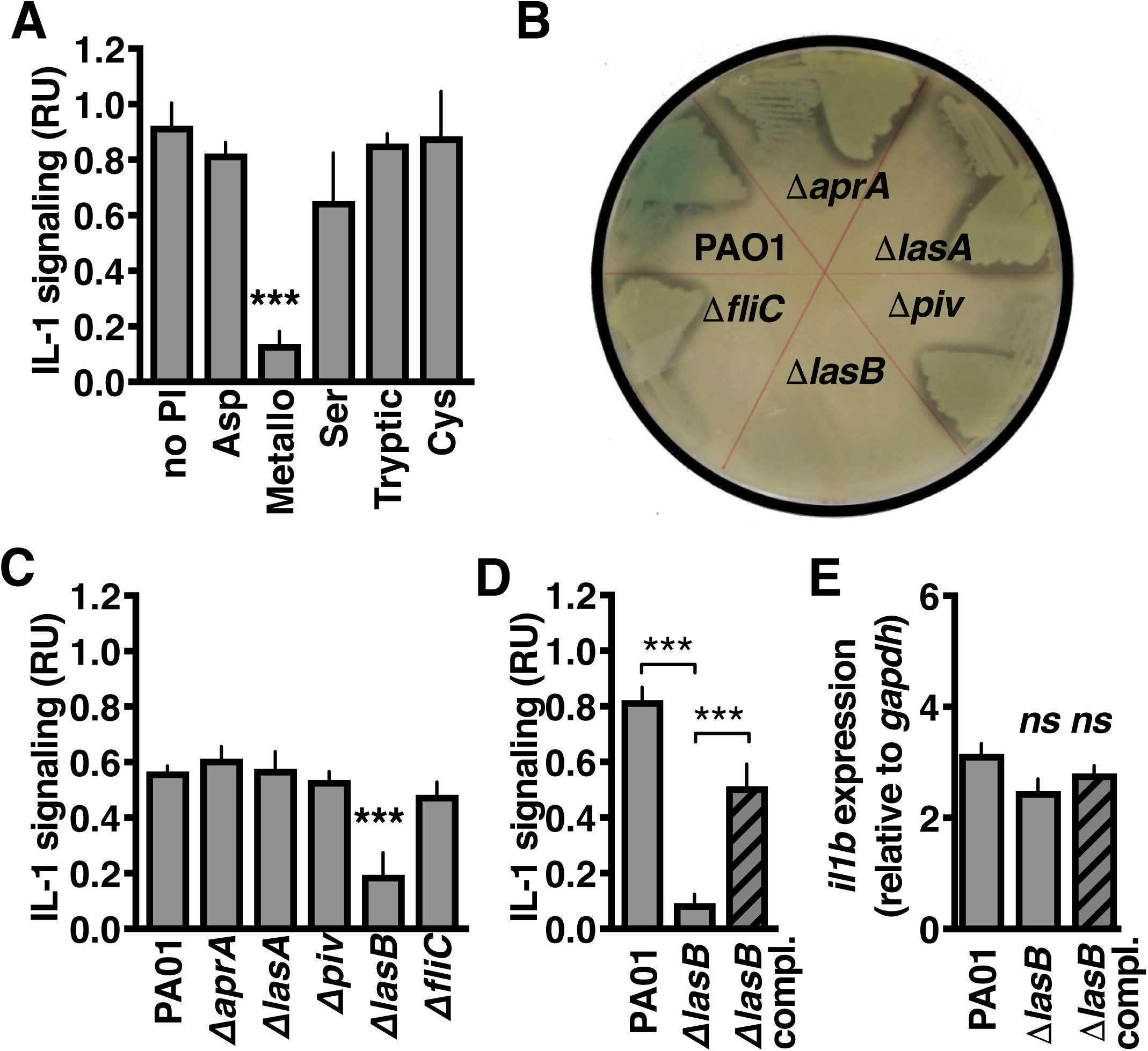
IL-1β is activated by the *P. aeruginosa* LasB protease. (**A**) Relative IL-1 signaling by caspase-1/11^-/-^ BMM 2 h post-infection by PAO1 that were previously incubated 1 h with the indicated protease inhibitors classes. (**B**) Visualization of bacterial proteolytic activity by decreased media opacity on LB agarose plates containing casein. (**C**) Relative IL-1 signaling by caspase-1/11^-/-^ BMM 2 h post-infection with isogenic mutant strains of PAO1. (**D**) Relative IL-1 signaling by caspase-1/11^-/-^ BMM 2 h post-infection by PAO1, Δ*lasB*, or plasmid-complemented Δ*lasB* and (**E**) *il1b* expression by real-time quantitative PCR. Where applicable, data are mean ± SEM and represent at least 3 independent experiments; significance determined by unpaired two-tailed Student’s T-test, \**P* < 0.05.

### LasB-activated IL-1β is active

Incubation with recombinant LasB was sufficient to convert recombinant human pro-IL-1β into an active form (Figure 4A). Further examination of pro-IL-1β cleavage by LasB, again using recombinant forms of each protein, showed several intermediate cleavage products which accumulate as a stable product that is degraded no further (Figure 4B), similar to what occurs upon IL-1β maturation by caspase-1 ^31^. Analysis of these fragments by Edman sequencing identified cleavage sites that were all in the N-terminus of pro-IL-1β. Examination of N-terminal truncated IL-1β by *in vitro* transcription/translation showed a defined region flanking the caspase-1 cleavage site (N-term fragment 117) is sufficient to generate active cytokine (Figure 4C). We further examined proteolysis in this ∼ 20 amino acid region with a series of internally quenched fluorescent peptides and found that LasB preferentially cleaved within the sequence HDAPVRSLN of pro-IL-1β (Figure 4D). Mass spectroscopy confirmed that LasB cleaved between Ser-121 and Leu-122 (Figure 4D, Figure S1), at a site conserved between mice and humans that matches the smallest IL-1β form we observed during SDS-PAGE (Figure 4B). This site also matches the substrate specificity profile for LasB (Figure 4E), which shows a distinct preference for cleaving peptide bonds when Ser or Thr are in the P1 position (amino-terminal side of bond) and hydrophobic amino acids such as Phe, Leu, Nle, Tyr, Trp and Ile in the P1′ position (Supplementary Spreadsheets 1-3), generated using a mass spectrometry-based substrate profiling assay previously validated with other microbial proteases ^32,33^. During infection, the signature of IL-1β-targeted proteolysis (Figure 4F) is consistent with a significant role for LasB-mediated maturation (hydrolysis of HDAPVRSLN) compared to caspase-1 (hydrolysis of EAYVHDAPV) ^30^. These data support the model that the pro-domain of IL-1β is promiscuous to protease activation and that the location of specific cleavages can dictate subsequent signaling activity.

**Figure 4.**
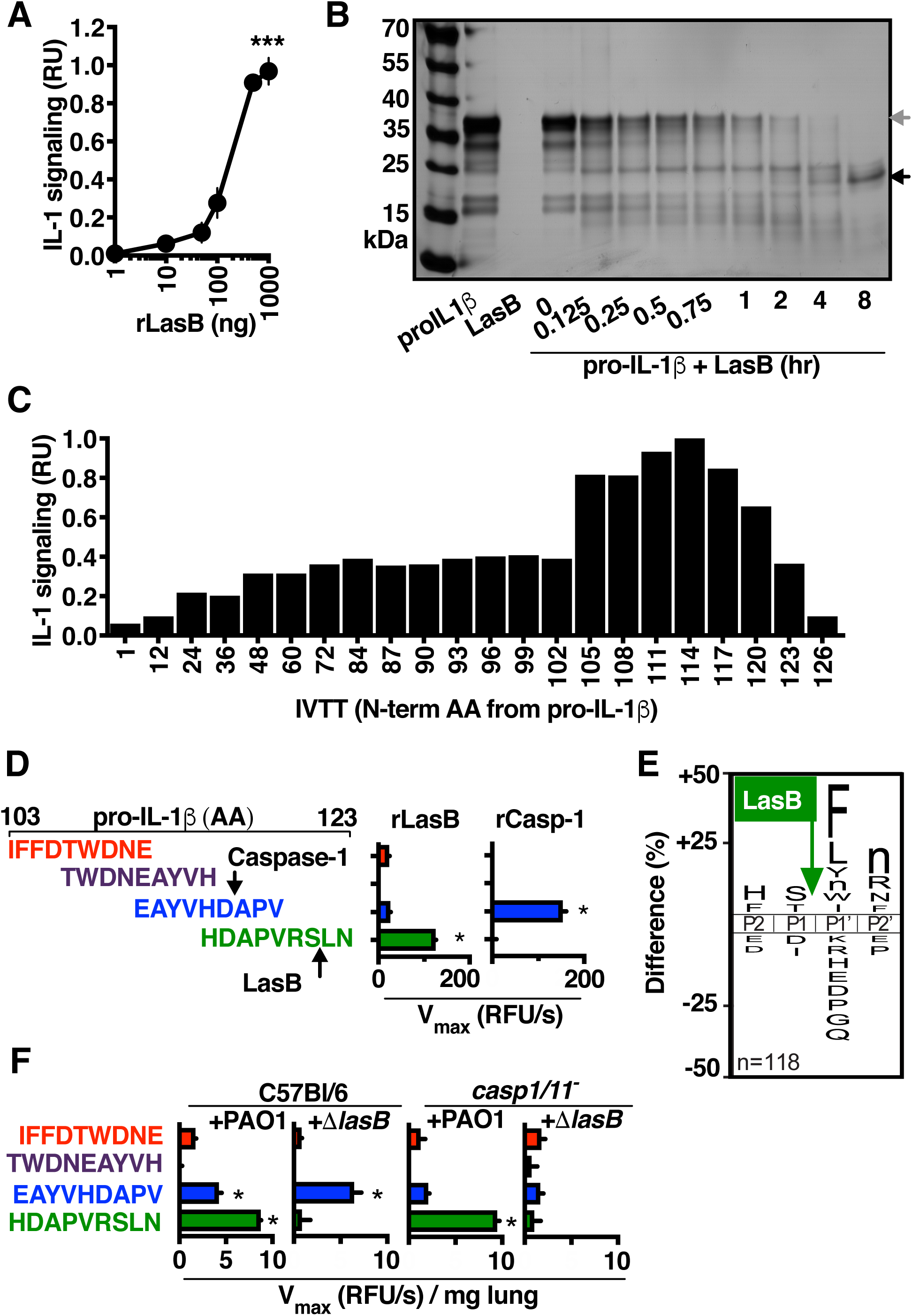
LasB-activated IL-1β is active. (**A**) IL-1 signaling activity by human pro-IL-1β after 2 h incubation with titrations of recombinant LasB. **(B**) SDS-PAGE analysis of recombinant human pro-IL-1β maturation by recombinant LasB. **(C**) Signaling activity of recombinant IL-1β N-terminal truncations generated using *in vitro* transcription/translation from each codon, 1 is full-length pro-IL-1β, 117 corresponds to the fragment generated by caspase-1 cleavage. **(D**) Cleavage of internally-quenched fluorescent IL-1β peptide fragments (amino acids 103-123 of human IL-1β) by recombinant LasB or caspase-1. (**E**) IceLogo frequency plot showing amino acids significantly enriched (above X-axis) and de-enriched (below X-axis) in the P2 to P2′ positions following incubation of LasB with a mixture of 228 tetradecapeptides. Cleavage occurs between P1 and P1′, lowercase “n” is norleucine. **(F**) Cleavage of internally-quenched fluorescent IL-1β peptide fragments by proteases within BAL collected from C57BL/6 or casp-1/11^-/-^ mice 24 h post-intratracheal infection with 10^7^ CFU of PAO1 or Δ*lasB.* Where applicable, data are mean ± SEM and represent at least 3 independent experiments; significance determined by unpaired two-tailed Student’s T-test, \**P* < 0.05.

### Metalloprotease inhibitors of LasB prevent IL-1β -mediated pathological inflammation

Since IL-1β inhibition protects against lung damage (Figure 1A, 1B), and because LasB drives IL-1β maturation (Figure 3C, 4D), we examined whether protease inhibitors active against LasB limit lung injury. Two investigational hydroxamate-based anti-neoplastic metalloprotease inhibitors, marimastat and ilomastat, inhibited LasB cleavage of the IL-1β-derived substrate (Figure 5A) and *P. aeruginosa* activation of IL-1β (Figure 5B) at sub-antimicrobial concentrations (Figure S2A). During murine pulmonary infection, marimastat and ilomastat each showed therapeutic effects to reduce IL-1β (Figure 5C), neutrophil recruitment (Figure 5D), pulmonary pathology (Figure 5E), and invasion (Figure S2B). Together this data suggests that inhibiting metalloproteases, including LasB, can reduce inflammation during infections by *P. aeruginosa*.

**Figure 5.**
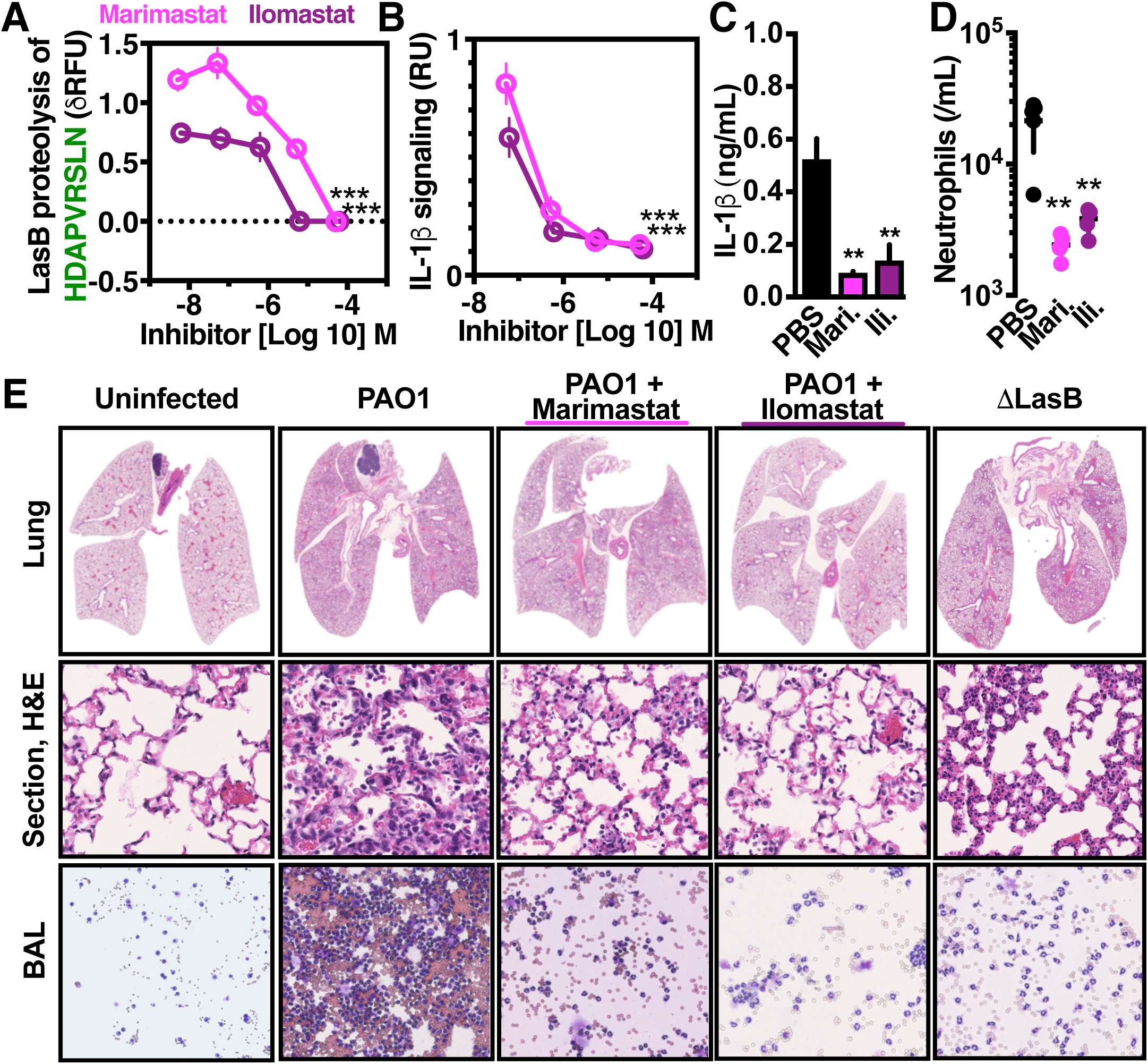
Metalloprotease inhibitors prevent pathological inflammation during *P. aeruginosa* infection. (**A**) Cleavage of internally quenched IL-1β fragment HDAPVRSLN by recombinant LasB incubated with titrations of Marimastat and Ilomastat. (**B**) IL-1 signaling by THP-1 macrophages 2 h post-infection with PAO1, MOI=10, incubated with titrations of Marimastat and Ilomastat. C57BL/6 mice intratracheally infected with 10^7^ CFU PAO1 and treated with 25 µg/kg Ilomastat, 25 µg/kg Marimastat, or PBS control. After 24 h, mouse BAL was harvested and (**C**) IL-1β measured by enzyme-linked immunosorbent assay and (**D**) neutrophils enumerated. (**E**) Representative histology sections cytological smears of bronchoalveolar lavage fluid prepared with differential MGG stain. Where applicable, data are mean ± SEM and represent at least 3 independent experiments. Significance determined by Mann-Whitney U-test (\**P* < 0.05, \*\**P* < 0.005).

## Discussion

Opportunistic *P. aeruginosa* lung infections can destroy tissue structure and impair organ function. Our findings reveal a mechanism by which a bacterial protease, LasB, contributes to pathological inflammation by directly activating IL-1β. LasB is one of the most abundant virulence factors in the lung microenvironment during *P. aeruginosa* infection and can cleave numerous host factors ^34^, even exerting broadly anti-inflammatory influences through destructive proteolysis of PAMPs such as flagellin ^35^, and various cytokines including IFN, IL-6, IL-8, MCP-1, TNF, trappin-2 and RANTES ^36-39^. Consequently, LasB-deficient bacteria may preferentially induce a KC, IL-6, and IL-8 dominant inflammatory response ^36^, whereas we find wild-type *P. aeruginosa* induce a strong IL-1β response.

LasB activates IL-1β through direct proteolytic removal of its inhibitory amino-terminal pro-domain, bypassing the necessity for host caspases. The LasB and caspase-1 mechanisms for generating mature IL-1β are distinguishable by substrate specificity (a hydrophobic P1’ *vs* aspartic acid P1 site), enzyme class (metalloprotease *vs* cysteine protease), and cellular source (microbial *vs* host). LasB activation of pro-IL-1β in both the intra- and extracellular milieu is entirely feasible, given the abundance of intracellular proteins released by pyroptosis and necrosis during infections ^10,40^ and the abundance of LasB ^41^. We recently hypothesized that IL-1β evolved as a sensor of diverse proteases ^30^, a model further supported by the present discovery of a *P. aeruginosa* protease with this activity.

In lung infection, LasB activation of IL-1β augments neutrophil recruitment and promotes destruction of the pulmonary tissue. IL-1β inhibition protects against this pathology, however, clinical interventions to date have utilized expensive biologics (e.g. IL-1R1 antagonists) associated with increased risk for severe infections ^30,42^. The proteolytic activation of IL-1β may be a more tractable pharmacological target, made possible by disambiguation of the molecular networks involved and, perhaps amenable to the repurposing existing proteases inhibitors. Alpha-1-antitrypsin suppresses NE-mediated degradation of the CF lung ^43,44^, potentially also limiting pro-IL-1β maturation by NE ^27^. This strategy may also act against pro-IL-1β maturation by LasB, which is also inhibited by alpha-1-antitrypsin ^45^. Metalloprotease inhibitors such as marimastat and ilomastat may also be beneficial in treating CF ^46^ not only for inhibiting matrix metalloproteases, but also by cross-inhibiting LasB (Figure 5).

## MATERIALS AND METHODS

### Bacterial strains and plasmids

All bacterial strains, plasmids, and primers used in this study are listed in Table 1. *lasB* and the upstream 260 bp regulatory region in PAO1 were cloned into pUC18T-mini-Tn7T-*hph* ^47^ using Polymerase Incomplete Primer Extension (PIPE) cloning ^48^ with primers lasB-F, lasB-R, Tn7-F, and Tn7-R. Transformants into Top10 cells were selected on LB agar plates containing 100 µg/mL Hygromycin B (Life Technologies). Stable complementation into PAO1 Δ*lasB* was performed as previously described ^47^, and transformants selected with 400 µg/mL Hygromycin B. pET-LasB with a C-terminal His tag was constructed by sequential PIPE cloning with the primers LasB-A, LasB-B, LasB-C, and LasB-D, and proteins were expressed and purified by conventional methods as previously described ^30^. pET-pro-IL-1β and the purification of pro-IL-1β have been previously described ^30^. Constructs for the expression of IL-1β mutants were generated by PIPE cloning from pET-pro-IL-1β ^30^ with the corresponding primers sets in Table 1, and proteins were expressed and purified in the same manner as for pro-IL-1β previously ^30^. Bacteria were routinely propagated in Luria broth (LB) medium at 37 °C. For infections, bacterial cultures were grown to late exponential phase (OD_600_ 1.2) then washed and diluted in PBS.

**TABLE 1.**
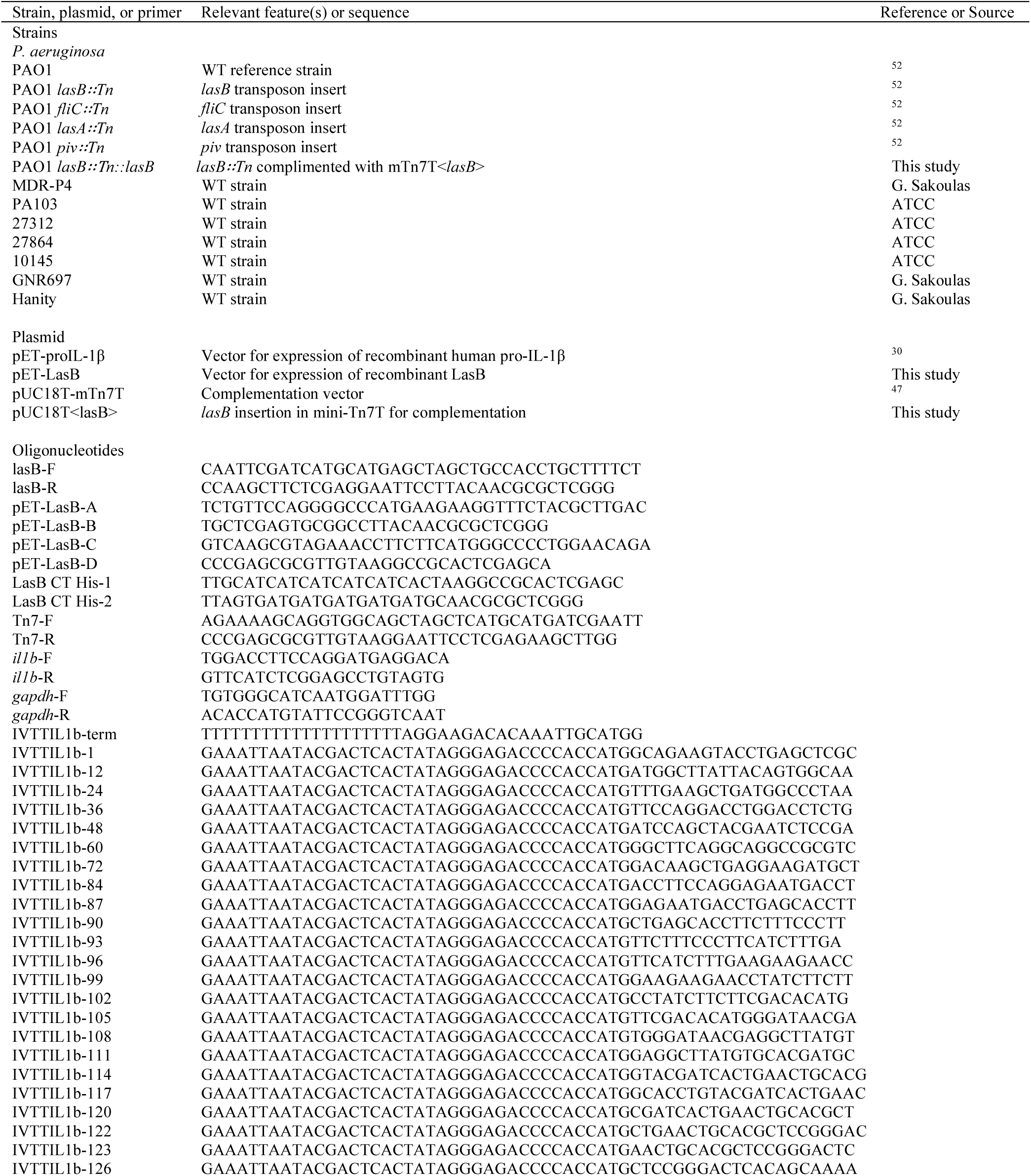
Bacterial strains, plasmids, and primers used in this study.

### Animal Experiments

The UCSD or Emory University Institutional Animal Care and Use Committees approved all animal use. Eight-to-ten week old male or female C57Bl/6 and isogenic caspase-1/-11^-/-^ mice were anesthetized with ketamine/xylazine intraperitoneally, then 10^7^ CFU PAO1 inoculated intratracheally in 30 µl of 1x PBS, 25 µg/kg Ilomastat, and 25 µg/kg Marimastat. Mice were euthanized by CO_2_ asphyxiation, and bronchiolar lavage fluid or lung homogenate were dilution plated onto LB agar plates for CFU enumeration, or quantification of cytokines or proteolysis. Bronchiolar lavage fluid cells were counted on a hemocytometer with cytologic examination on cytospin preparations fixed and stained using Hema 3 (Fisher HealthCare™). Histologic sections were prepared from formalin-fixed and paraffin-embedded lungs, stained with hematoxylin and eosin (H&E). Cytospin and histology slides were imaged on a Hamamatsu Nanozoomer 2.0Ht Slide Scanner.

### *In vitro* infection models

Macrophages were generated from femur exudates of wild-type C57Bl/6 (Jackson Laboratories) or caspase-1/11^-/-^ (kindly provided by R. Flavell) mice using M-CSF containing L929 cell supernatants as previously ^30^. THP-1, HL60, and A549 cells were propagated by standard protocols detailed previously ^49^. One hour before infection, the media was replaced with RPMI lacking phenol red, fetal bovine serum, and antibiotics. Inhibitor treatments were added 1 h before infection and include: 20 µg/mL Anakinra (Amgen), 100 ng/mL rIL-1β (R&D Systems), 5 µM caspase inhibitors zVAD-fmk, YVAD-fmk, DEVD-fmk, and IETD-fmk (R&D Systems), 10 µg/mL complete protease inhibitor cocktail (Roche), 1x protease inhibitors AEBSF, Antipain, Aprotinin, Bestatin, EDTA, E-64, Phosphoramidon, Pepstatin, and PMSF (G-Biosciences). Except when noted, cells were routinely infected by co-incubation with *P. aeruginosa* at a multiplicity of infection of 10, spun into contact for 3 min at 300 g, and cells or supernatants were harvested for analysis after 2 h.

### Cytokine measurements

Relative IL-1 signaling by cells was measured in 50 µl of supernatant from infected or treated cells, then incubated with 1 µM okadaic acid 30 min before transfer onto transgenic IL-1R reporter cells (Invivogen). After 18 h, reporter cell supernatants were analyzed for secreted alkaline phosphatase activity using HEK-Blue Detection reagent (Invivogen). Cytokines were quantified by enzyme-linked immunosorbent assay following the manufacturer’s protocol (R&D Systems). Expression was examined in cells lysed with RIPA (Millipore). RNA was isolated (Qiagen), cDNA synthesized with SuperScript III and Oligo(dT)20 primers (Invitrogen), and qPCR performed with KAPA SYBR Fast (Kapa Biosystems) with primers for *il1b* and relative expression normalized to *gapdh* and compared by ΔΔCt as previously ^50^. *In vitro* transcription/translation was performed with the corresponding primers in **Table 1** using pET-pro-IL-1β as a template and following the manufacturer’s recommendations in 10 µl reaction volumes (TNT Coupled Reticulocyte Lysate; Promega). Loading for IL-1R reporter assays was normalized by total IL-1β product measured by enzyme-linked immunosorbent assay (R&D Systems).

### Substrate specificity profiling

10 nM LasB was incubated in triplicate with a mixture of 228 synthetic tetradecapeptides (0.5 µM each) in PBS, 2mM DTT as described previously ^51^. After 15, 60, 240 and 1200 min, aliquots were removed, quenched with 6.4 M GuHCl, immediately frozen at -80°C. Controls were performed with LasB treated with GuHCl prior to peptide exposure. Samples were acidified to pH<3.0 with 1% formic acid, desalted with C18 LTS tips (Rainin), and injected into a Q-Exactive Mass Spectrometer (Thermo) equipped with an Ultimate 3000 HPLC. Peptides separated by reverse phase chromatography on a C18 column (1.7 µm bead size, 75 µm x 20 cm, 65°C) at a flow rate of 400 nl/min using a linear gradient from 5% to 30% B, with solvent A: 0.1% formic acid in water and solvent B: 0.1% formic acid in acetonitrile. Survey scans were recorded over a 150–2000 m/z range (70000 resolutions at 200 m/z, AGC target 1×10^6^, 75 ms maximum). MS/MS was performed in data-dependent acquisition mode with HCD fragmentation (30 normalized collision energy) on the 10 most intense precursor ions (17500 resolutions at 200 m/z, AGC target 5×10^4^, 120 ms maximum, dynamic exclusion 15 s).

Peak integration and data analysis were performed using Peaks software (Bioinformatics Solutions Inc.). MS^2^ data were searched against the tetradecapeptide library sequences and a decoy search was conducted with sequences in reverse order with no protease digestion specified. Data were filtered to 1% peptide and protein level false discovery rates with the target-decoy strategy. Peptides were quantified with label free quantification and data normalized by LOWESS and filtered by 0.3 peptide quality. Missing and zero values are imputed with random normally distributed numbers in the range of the average of smallest 5% of the data±SD. Enzymatic progress curves of each unique peptide were obtained by performing nonlinear least-squares regression on their peak areas in the MS precursor scans using the first-order enzymatic kinetics model: Y = (plateau-Y_0_)×(1-exp(–t×k_cat_/K_M_×[E_0_]))+Y_0_, where E_0_ is the total enzyme concentration. Nonlinear regression was performed on cleavage products only if the following criteria were met: Peptides were detected in at least 2 of the 3 replicates and the peak intensity of peptides increased by >50,000 and >5-fold over the course of the assay. Proteolytic efficiency was solved from the progress curves by estimating total enzyme concentration and is reported as k_cat_/K_M_ and clustered into 8 groups by Jenks optimization method. IceLogo software was used for visualization of amino-acid frequency using cleavage sequences in the top 3 clusters (118 most efficiently cleaved peptides). Mass spectrometry deposited:ftp://massive.ucsd.edu/MSV000081623.

### Protease Measurements

Internally-quenched peptides 7-Methoxycoumarin- (Mca) labeled on the amino terminus and 2, 4-dinitrophenyl (Dnp) on the carboxy terminus were synthesized with the sequences of IFFDTWDNE, TWDNEAYVH, EAYVHDAPV, and HDAPVRSLN, corresponding to amino acids 103-111, 107-115, 111-119, and 115-123 of the reference human pro-IL-1β sequence (UniProt: P01584; CPC Scientific). In triplicate, 10 µM peptides were incubated in PBS, 1 mM CaCl_2_, 0.01% Tween-20, with 5 nM human caspase-1 (Enzo) or LasB (Elastin Products Co.). The reaction was continuously monitored using an EnSpire plate reader (PerkinElmer) with 323nm fluorophore excitation and 398nm emission and the maximum kinetic velocity calculated as previously ^30^. The cleavage site was determined by incubating 10 nM of LasB with 10 µM of HDAPVRSLN. At 20, 40 and 60 min intervals each reaction was quenched with 6.4 M GuHCl and the cleavage products desalted and analyzed by mass spectrometry as described above, except using a 20-min linear gradient from 5% to 50% B and only selecting top 5 peptides for MS/MS.

### Statistical analysis

Statistical significance was calculated by unpaired Student t test (*, P < 0.05; **, P < 0.005) using GraphPad Prism unless otherwise indicated. Data are representative of at least three independent experiments. For iceLogo plots only amino acids with significantly (P < 0.05) increased or decreased frequency are shown.

## Acknowledgments

We thank Christopher Lietz (UCSD) for assistance with mass spectrometry and data analysis, the UCSD Histopathology Core facility, the UCSD Neuroscience Microscopy Shared Facility (P30 NS047101), Jason Munguia (UCSD) for technical assistance, Colin Manoil (UW) for *Pseudomonas* transposon mutants (P30 DK089507) and Joanna Goldberg (Emory) for helpful advice and discussions. J.S. received support from NIH/NIGMS T32 GM007752, E.A.S. from NIH/NCI T32 CA121938, J.M.K. from a UC President’s Postdoctoral Fellowship, Z.J. from the UC San Diego Chancellor’s Research Excellence Scholarship, A.J.O. from NIH/NIBIB AI1333393, V.N. from NIH/NICHD grant U54 HD090259 and NIH/NHLBI R01 HL125352, and C.L. the A.P. Giannini Foundation and NIH/NIAID K22 AI130223. C.N.L. has a research agreement with Antabio examining inhibitors of LasB. The content is solely the responsibility of the authors and does not necessarily represent the official views of the National Institutes of Health.

## Author contributions

J.S., A.J.O., V.N., and C.N.L. designed experiments and interpreted the data. J.S., D.L., J.K., J.O., Z.J., E.A.S., A.J.O., and C.N.L conducted the studies. J.S., V.N., and C.N.L. wrote the manuscript with the assistance of all of the authors.

